# Global patterns and evolution of avian song complexity

**DOI:** 10.64898/2026.08.04.742837

**Authors:** Martin Bulla, Peter Mikula, Wolfgang Forstmeier, Tereza Petrusková, Tomáš Albrecht

## Abstract

Why birds evolved complex songs has been a long-standing puzzle. Here, by quantifying song complexity in 4,940 species (83% of all passerines), we show a latitudinal gradient in song complexity, with the most complex songs occurring in temperate, open-habitat bird assemblages characterized by relatively inconspicuous plumage. Using comparative analyses controlled for common ancestry, we find little support for traditional explanations based on sexual selection intensity, intelligence signaling, or species richness. Instead, complex songs evolve primarily in song-learning passerines, especially migrants, habitat generalists, and species breeding in open habitats where structured acoustic signals transmit more efficiently. Together, our findings indicate that global variation in song complexity emerges from the interaction between vocal learning, ecological context, and evolutionary history rather than from a single selective driver.

## Main Text

Many organisms use acoustic signals to attract mates and repel rivals (*1, 2*). Among the most spectacular and enigmatic examples are the elaborate songs of passerine birds (order: Passeriformes). Passerine songs vary from single notes to repertoires with hundreds of distinct elements, exemplified by iconic mockingbirds and nightingales (*3, 4*). Numerous hypotheses have been proposed to explain between-species differences in bird song complexity, such as sexual selection, species recognition, cognition, habitat structure, and trade-offs with other signals such as plumage coloration, but empirical support remains mixed (*3–10*). Yet, despite extensive work on avian visual signals and their global evolutionary patterns (*11, 12*), we lack even a basic description of how song complexity distributes across the globe, and thus a comprehensive test of the main competing hypotheses across all passerines.

Here, (i) to provide a global map of song complexity and (ii) to test the most influential hypotheses explaining interspecific variability in song complexity, we quantified male song complexity for 4,940 passerine species, representing ∼83% of extant species (*13*), using 18,065 song recordings (Fig. 1) from large citizen-science databases (Xeno-canto https://xeno-canto.org/ and Macaulay Library https://www.macaulaylibrary.org/). We predict strong clade-specific effects because song acquisition differs fundamentally between Suboscines with innate song and song-learning Oscines (*14*).

**Fig. 1.**
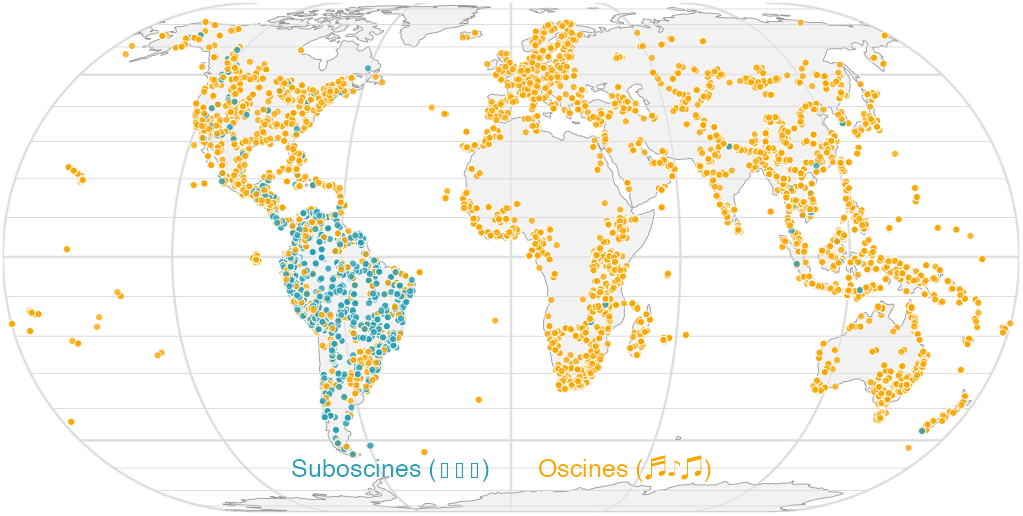
Global distribution of 18,488 analyzed passerine song recordings. Each point represents one recording. Color denotes clade: Suboscines (largely innate song; blue) and Oscines (song learners; orange).

We defined song complexity as the number of different element types in a sequence of 50 song elements (for details, see (*4*)). Our measure correlates positively with published metrics of song complexity (Fig. S1) (*4*), such as syllable repertoire size (r = 0.77; N = 129 species), number of unique syllables per song (r = 0.53; N = 226), and song repertoire size (r = 0.62; N = 182), as well as with size of HVC (*15*), an avian brain region involved in song production and vocal learning (r = 0.58; N = 57).

## Results and discussion

Song complexity follows a latitudinal gradient. The most complex songs are produced by species assemblages with relatively low male colorfulness, a measure of plumage color diversity, in temperate regions of the Northern Hemisphere, particularly open habitats in the Palearctic and Saharo-Arabian zoogeographic regions and parts of the Nearctic region (Fig. 2A and S2) (*11*). In contrast, songs are relatively simple in the tropics, where species assemblages tend to have higher male-plumage colorfulness (Fig. 2A and S2) (*11*). Although these associations may partly reflect the non-random spatial distribution of passerine lineages rather than adaptive responses per se, cross-species analyses controlled for common ancestry confirm a latitudinal gradient in song complexity (see univariate outputs in Fig. S3, Table S1).

**Fig. 2.**
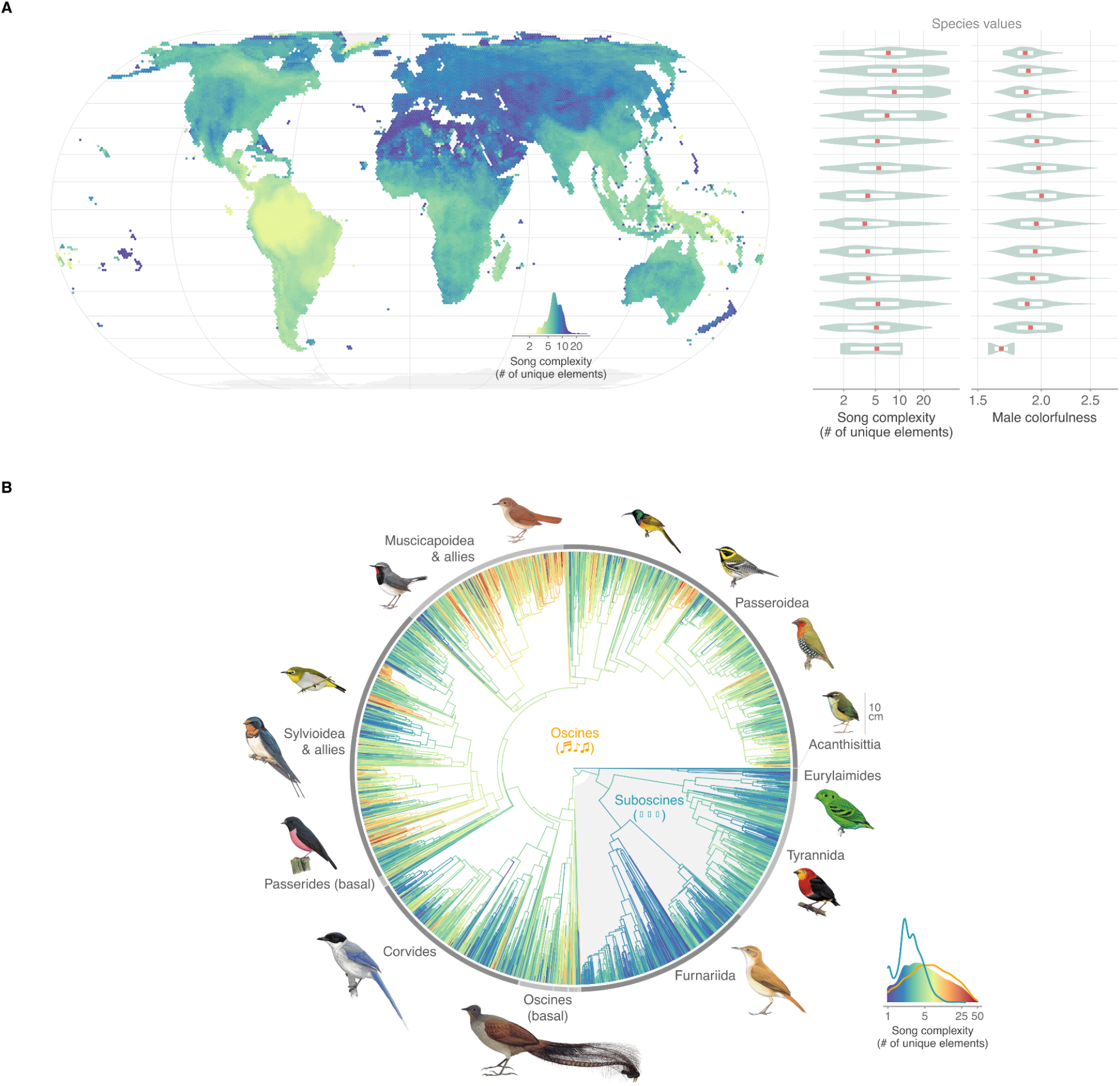
Global distribution and evolution of song complexity across passerines. (**A**) Geographic distribution of song complexity across species assemblages, defined for 1° grid cells (∼ 100 × 100 km) based on breeding ranges. Color encodes assemblage-mean song complexity, with the color scale indicating both value mapping and distribution (kernel density; n = 14,940 cells). Colors are clipped at the 2.5th and 97.5th percentiles for visualization. Violin plots show species-level distributions of song complexity and male colorfulness across 10° latitudinal bands (species assigned by breeding-range centroid). Red dots represent medians; white boxes show the 25th and 75th percentiles. (**B**) Distribution of song complexity across a maximum credibility phylogeny (based on 100 trees from http://birdtree.org). Color encodes observed species-mean song complexity at the tips (n = 4,940) and maximum-likelihood ancestral-state estimates at internal nodes, with the color scale indicating both value mapping and distribution (kernel density) and overlaid curves showing clade-specific distributions (orange: Oscines, blue: Suboscines). Highlighted are 10 major groups of passerines, represented by 14 illustrative species, roughly scaled according to size, except for the downscaled representative of the basal Oscines (should be three times larger); starting with Acanthisittia and going clockwise, the pictures depict *Xenicus gilviventris* (10 cm body size), *Calyptomena viridis* (16 cm; Eurylaimides), *Pipra aureola* (11 cm; Tyrannida), *Furnarius rufus* (19.5 cm; Furnariida), *Menura novaehollandiae* (103 cm; Oscines (basal)), *Cyanopica cyanus* (35 cm; Corvides), *Petroica rodinogaster* (12 cm; Passerides (basal)), *Hirundo rustica* (18 cm; Sylvioidea & allies), *Zosterops poliogastrus* (12 cm; Sylvioidea & allies), *Luscinia pectorali*s (15 cm; Muscicapoidea & allies), *Luscinia megarhynchos* (17 cm; Muscicapoidea & allies), *Nectarinia violacea* (16 cm; Passeroidea), *Dendroica townsendi* (12 cm; Passeroidea) and *Mandingoa nitidula* (11 cm; Passeroidea). Species names follow the taxonomy of the BirdTree phylogenetic backbone used in the analyses. Illustrations sourced from Birds of the World, Cornell Lab of Ornithology.

We find a strong evolutionary signal in song complexity across passerines, as well as within their two major clades, Oscines and Suboscines (Pagel’s λ = 0.74, 0.71, 0.68, respectively; Fig. 2B). This phylogenetic effect is unevenly distributed across taxonomic levels. Clade explains 13% of the phenotypic variance in song complexity (Table S2), with song-learning Oscines producing on average substantially more complex songs (estimate [95% CI] = 0.87 [0.42–1.32] standard deviations) than Suboscines with innate song (Fig. 2B, Table S3). Yet, many Oscine lineages also produce relatively simple songs (Fig. 2B and S4; e.g., *Locustella* warblers, *Emberiza* buntings and tits), suggesting that vocal learning is a necessary but not sufficient condition for the evolution of highly complex songs. Apart from clade, taxonomic levels above family explain little phenotypic variance (6%), while substantial variance at family (13%) and genus (24%) levels indicates that song complexity can diverge relatively rapidly (Fig. 2B; Table S2).

Three main hypotheses have been proposed to explain the evolution of song complexity.

### (1) ‘Intensity of sexual selection’ hypothesis

Strong intra- and inter-sexual selection has been proposed to favor males with complex songs (*16–18*). However, we find no strong evidence that sexual selection predicts variation in passerine song complexity (Fig. 3), based on multiple traditional proxies for sexual selection intensity: sexual size dimorphism (*19, 20*), sexual plumage-color dimorphism (*12*), male-like coloration, reflecting similarity between female and male plumage (*6, 11, 12*), degree of social polygyny (*4, 7*), territoriality (i.e., the intensity of sexual selection on males) (*21, 22*), and stability of social bonds (i.e., opportunity for mate choice) (*21, 22*). Consistent with this result, song complexity is also unrelated to the level of extra-pair paternity (Table S4). A partial exception is observed in Suboscines, where territorial species show a noisy trend toward lower song complexity than non-territorial species (Fig. 3 and S3; see also (*23*)).

**Fig. 3.**
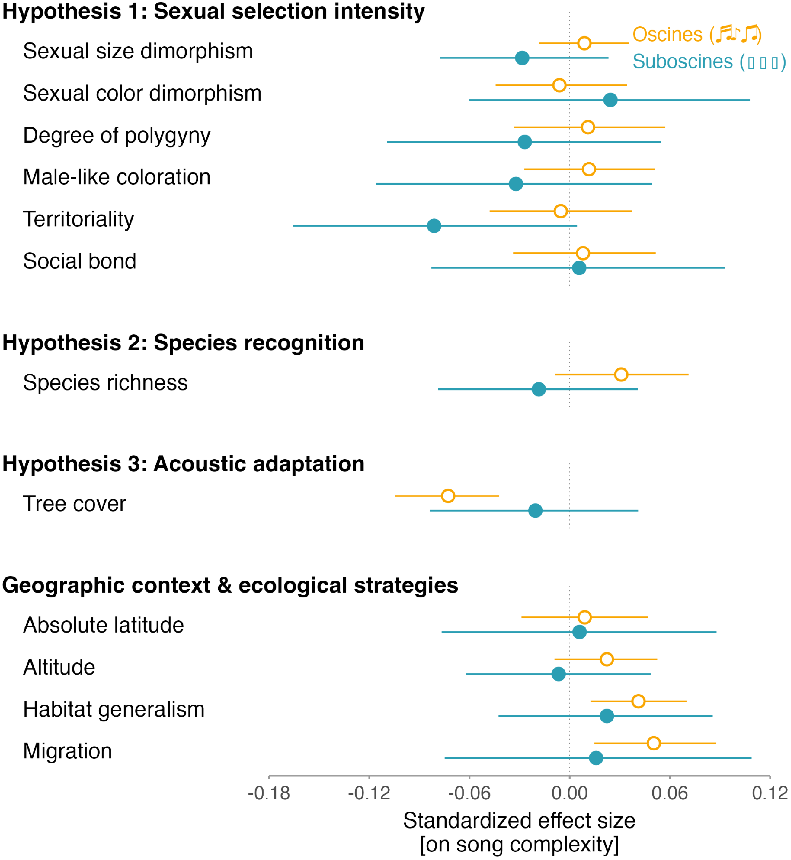
Predictors of song complexity. Dots with horizontal lines represent estimated standardized effect sizes and their 95% credible intervals from a multivariate mixed-effect taxonomic model containing all predictors and their interactions with clade (Table S3). Color and dot type indicate clade: orange open dots represent song-learning Oscines, blue filled dots Suboscines with largely innate song. The estimate for clade alone is available in Table S3. Alternative analyses, including clade-specific models, a model accounting for heteroscedasticity, a model using male colorfulness instead of male-like coloration, a phylogenetic regression (PGLS), and univariate models, yielded results consistent with those from the model depicted here (Fig. S3 and S6, Table S1). Species *N* = 4,386; 3,329 Oscines, 1,057 Suboscines.

### (2) ‘Species recognition’ hypothesis

Reduced song complexity has been predicted in areas with high numbers of breeding passerine species, since overly complex songs may hinder rapid and efficient species recognition (*24, 25*). However, we find no support for such a negative association between species richness and song complexity (Fig. 3 and S3).

### (3) ‘Acoustic adaptation’ hypothesis

Dense habitats are expected to select for structurally simple acoustic signals because sounds degrade during transmission via absorption, reverberation, and scattering, processes typically stronger in closed, forested habitats (*26–29*). Indeed, in song-learning Oscines, song complexity is lower in forests than in open habitats (Fig. 3 and S3), consistently across latitudes (i.e., not restricted to tropical or temperate forests; Table S5). This tree cover effect was not observed for peak song frequency (*30*).

The habitat axis that predicts song complexity also structures visual signals: male colorfulness increases with forest-dependent lifestyle and toward lower latitudes (*11*). We therefore examined whether the association between song complexity and male colorfulness depends on ecological context. In Oscines, song complexity was negatively associated with male colorfulness only in temperate forests; this association was absent in tropical forests, open habitats, and Suboscines (Fig. 4, Table S6). Thus, the apparent negative song-color relationship (Fig. 2A) is not general. Why such context-dependent partitioning between acoustic and visual signals appears most evident in temperate forests remains unclear, perhaps because acoustic constraints, though present, are less extreme than in tropical forests (*31, 32*).

**Fig. 4.**
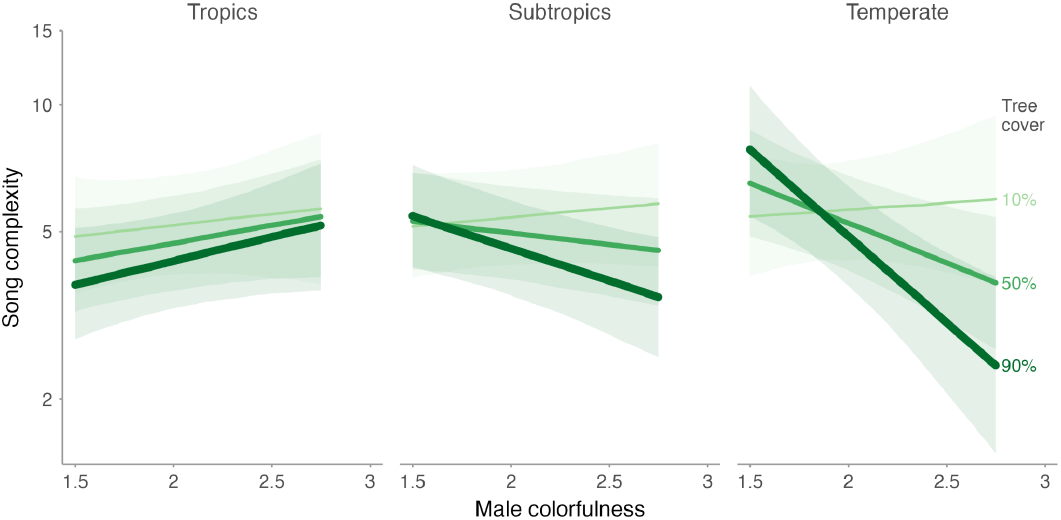
Relationship between song complexity, male colorfulness, tree cover, and latitude for song-learning Oscines. Predicted relationships (solid lines) and 95% confidence intervals (shaded ribbons), derived from a linear mixed-effects model containing the four-way interaction of clade, male colorfulness, tree cover and latitude, while controlling for taxonomy (Table S6). Color and line width indicate tree cover: 10% (light green, thin line), 50% (medium green, medium line), and 90% (dark green, thick line). Panels represent distinct latitudinal contexts: Tropics (proxied by 0° latitude), Subtropics (23°), and Temperate (45°) regions. Song complexity (number of element types) is plotted on a log-scale, with y-axis tick labels back-transformed to original units.

### Geographic context and ecological strategies

We examined geographic context and ecological strategies to describe, in general terms, which species have evolved complex songs. In a multivariate model, song complexity is unrelated to absolute latitude and altitude; in Oscines, however, song complexity is higher in migratory species and habitat generalists (Fig. 3, Fig. 5A, Table S1), indicating that these ecological strategies largely account for the observed latitudinal pattern. Their associations with song complexity are additive, with no evidence for an interaction between migration and habitat generalism (Fig. S3, Fig. S5, Table S7).

**Fig. 5.**
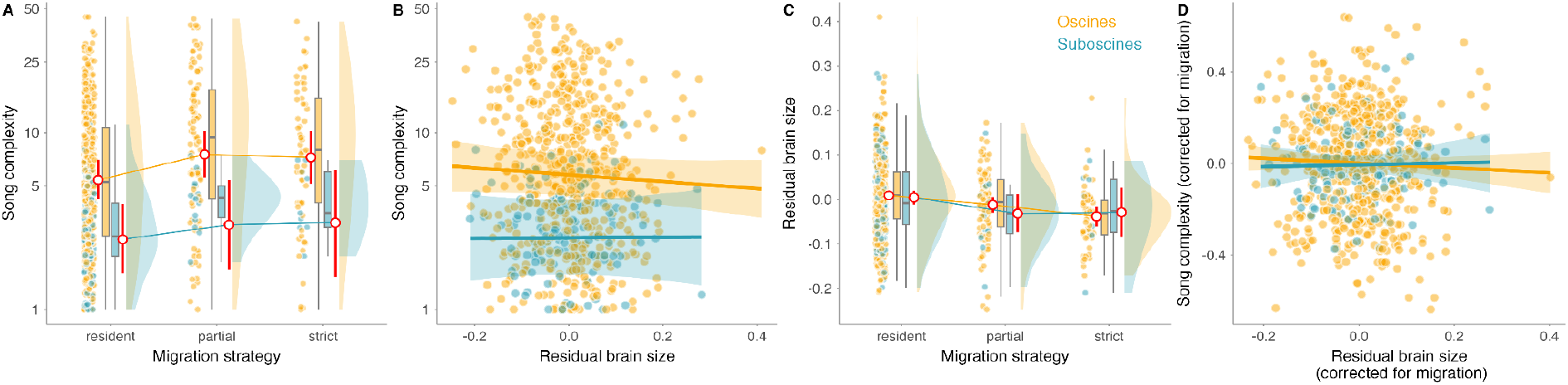
Clade specific relationships between song complexity, migratory strategy and brain size. **(A)** Song complexity is higher in migratory species. **(B)** Song complexity shows a weak negative relationship with residual brain size in Oscines. **(C)** Migratory species tend to have smaller residual brain sizes than resident species. **(D)** After correcting both song complexity and residual brain size for migratory strategy, their relationship is statistically unclear. The apparent reduction of the Oscine-Suboscine difference in D is expected because both variables are residuals from mixed models that remove taxonomic structure; thus, panel D shows the within-taxonomic, migration-corrected association rather than raw clade differences. Colors indicate clade: orange = song-learning Oscines; blue = Suboscines with largely innate song. Points represent species-level data, jittered to improve visibility; N = 679 species with brain-size data. Red points and vertical bars in **A** and **C**, and fitted lines with shaded areas in **B** and **D**, show predictions and 95% confidence intervals from univariate mixed-effects models used for visualization. Models included clade interactions and nested taxonomic random intercepts for genus, family, superfamily, parvorder and infraorder. Boxplots show medians, interquartile ranges and whiskers extending to 1.5 times the interquartile range or to the minimum/maximum value, whichever is smaller. Song complexity in **A** and **B** is plotted on a log scale, with tick labels back-transformed to original units.

A popular interpretation of such patterns has been that species living in variable environments have evolved complex songs as an honest ‘signal of intelligence’, often proxied by brain size (*33*), thereby facilitating female choice of an optimal mate (*5, 10, 16, 17*). Despite conceptual concerns that courtship traits evolve primarily for the benefit of the signaler rather than the receiver (*34*), the hypothesis of honest signaling of intelligence remains widely accepted (*16, 35–37*).

Contrary to expectations, song complexity is negatively correlated with brain size (Fig. 5B), either measured as absolute size or after accounting for differences in body mass (Table S8). We consider the negative relationship to be largely a spurious byproduct of variation in migratory behavior. Migrants tend to have smaller brains (Fig. 5C, Table S9) (*38*) yet more complex songs than residents (Fig. 3, Fig. 5A, Table S1) (*8*). After accounting for migratory behavior, the association between song complexity and brain size vanishes, resulting in a statistically unclear negative trend (Fig. 5D, Table S10). This lack of a positive relationship between song complexity and brain size remains inconsistent with the idea that complex songs evolved through adaptive female choice as an honest signal of brain size or intelligence.

## Conclusions

Our study provides the first global synthesis of passerine song complexity and challenges several long-standing explanations for its evolution. Despite pronounced geographic and phylogenetic structure, song complexity is largely unrelated to proxies of sexual selection and brain size, providing little support for the widespread view that elaborate songs primarily function as ‘honest’ sexually selected signals of cognitive ability. Instead, patterns of song complexity are most strongly associated with clade-specific constraints, habitat structure, and species ecology, with effects largely restricted to song-learning Oscines. Our results further indicate that trade-offs between acoustic and visual signals are strongly context dependent rather than general features of avian communication. These findings suggest that extreme song complexity arises not from a single selective driver but from interactions among vocal learning, ecological context, and evolutionary history, emphasizing the need to reconsider simple adaptive narratives of acoustic signal evolution.

### Methods and supporting materials

An interactive HTML document gives detailed methods, supplementary figures and tables, along with R-code used to generate all display items, and links to data https://martinbulla.github.io/passerine_song_complexity_v2/versions/v2.0.2/ (*39*).

## Acknowledgements

We thank all contributors and administrators of the Xeno-canto and Macaulay library platforms – without their efforts this study would not have been possible. We are grateful to Barbora Blažková for help with collecting the recordings, Henrik Brumm for help with designing the complexity metric, Mihai Valcu, Fränzi Korner-Nievergelt and Liam J. Revell for advice on the statistics, members of the Department of Ornithology, Max Planck Institute for Biological Intelligence (especially Mihai Valcu and Bart Kempenaers) and Henrik Brumm for feedback and suggestions. Adam, Anička, Barbora, Jana, Julie, Jonáš, and Majlen for patience and support.

## Author contributions

T.A. and P.M. conceived the study. T.A. supervised the study. All authors developed the methods. P.M. collected song complexity data, P.M. & M.B. collected data on song complexity predictors. M.B. performed formal analyses with input from W.F., T.A. & P.M. and prepared the Supplementary Information. T.A., M.B., W.F., P.M. (alphabetically) wrote the first draft, and all authors contributed to its revision.

